# Decomposing Simon task BOLD activation using a drift-diffusion model framework

**DOI:** 10.1101/809947

**Authors:** James R McIntosh, Paul Sajda

## Abstract

The Simon effect is observed in spatial conflict tasks where the response time of subjects is increased if stimuli are presented in a lateralized manner so that they are incongruous with the response information that they represent symbolically. Previous studies have used fMRI to investigate this phenomenon, and while some have been driven by considerations of an underlying model, none have attempted to directly tie model and BOLD response together. It is likely that this is due to Simon models having been predominantly descriptive of the phenomenon rather than capturing the full spectrum of behavior at the level of individual subjects. Sequential sampling models (SSM) which capture full response distributions for correct and incorrect responses have recently been extended to capture conflict tasks.

In this study we use our freely available framework for fitting and comparing non-standard SSMs to fit the Simon effect SSM (SE-SSM) to behavioral data. This model extension includes specific estimates of automatic response bias and a conflict counteraction parameter to individual subject behavioral data. We apply this approach in order to investigate whether our task specific model parameters have a correlate in BOLD response. Under the assumption that the SE-SSM reflects aspects of neural processing in this task, we go on to examine the BOLD correlates with the within trial expected decision-variable. We find that the SE-SSM captures the behavioral data and that our two conflict specific model parameters have clear across subject BOLD correlates, while other model parameters, as well as more standard behavioral measures do not. We also find that examining BOLD in terms of the expected decision-variable leads to a specific pattern of activation that would not be otherwise possible to extract.

## Introduction

It is somewhat surprising that when a stimulus is presented at a location irrelevant to the required response, that this location should feed into the decision process. This is what spatial conflict, and specifically the Simon effect^1, 2^ demonstrates: the average response time (RT) of subjects is increased, and accuracy decreased if stimuli are presented in a lateralized manner so that they are incongruous with the response information that they represent symbolically.

The Simon effect deviates from the behavior that would be predicted by a simple application of the drift-diffusion model (DDM)^3–5^. This deviation appears due to subjects making use of task-irrelevant information, and manifests in the shape of subjects’ RT distributions^6–9^. In recent years sequential sampling models (SSM) for conflict^10, 11^) and the Simon task specifically^12^ have been developed which fully capture RT distributions and accuracy.

The explained operation of the Simon effect SSM (SE-SSM^12^) within a conflict trial is rooted in models of cognitive control and prior models of the Simon effect^13, 14^. The SE-SSM, which we make use of here extends the SSM in two specific ways. It introduces a conflict counteraction term, which acts to boost the decision-variable in conflict trials; and it also adds an initial bias to the decision-variable, which more traditionally would be thought of as activation of an automatic^15^ pathway. From the perspective of model fitting, both of these parameters are needed. This is because while a bias term captures the increased mean response time in incongruent trials, the conflict counteraction term is needed to capture a compensatory decrease in the standard deviation of the RT. In previous work the SE-SSM was shown to explain behavior during a Simon task, and predict changes in RT induced by transcranial random noise stimulation (tRNS).

Such models lend themselves to enhance our understanding of the Simon task^16^ and enable new analysis in neuroimaging studies^17^. Previous studies have used fMRI to investigate the Simon effect^18–23^, and while some of these have used analyses derived from Simon effect model considerations, none of these models could account for full RT and accuracy distributions. These previous analyses have however highlighted potential regions of interest such as dorsolateral prefrontal cortex (DLPFC), pre-supplementary motor area (pre-SMA), and anterior cingulate cortex^17^ (ACC). The power of the SSM is not only that it can model behavior but that it is cast at a level of abstraction where its specific components are directly interpretable under the assumption that it captures some core principles of decision making used in the brain.

In this work we develop a model fitting framework which can be used to fit non-standard SSMs. We use this framework to fit the SE-SSM and show that it captures response time and accuracy of subjects that performed a Simon task. We go onto make use of the model fits by directly relating its parameters to fMRI data, and finally we exploit the high-temporal resolution of the model to decompose the BOLD signal.

## Methods

### Data source and task

Data analyzed in this manuscript was obtained from the OpenfMRI database under an ODC Public Domain Dedication and Licence (PDDL). Its accession number is ds000101^24^. We reproduce here relevant details of the data, and refer the reader to the OpenfMRI database for a full description. Healthy adults (n = 21, male: 12, female: 9, age mean: 30, age std: 7) were scanned in a Siemens Allegra 3.0 T scanner, with a standard Siemens head coil, located at the NYU Center for Brain Imaging. 151 contiguous echo planar imaging (EPI) whole-brain functional volumes were obtained (TR = 2000ms; TE = 30ms; flip angle= 80, 40 slices, matrix= 64 × 64; FOV= 192mm; acquisition voxel size= 3 × 3 × 4mm^3^) during each of the two sessions. A high-resolution T1-weighted anatomical image was also acquired using a magnetization prepared gradient echo sequence (MPRAGE, TR = 2500ms; TE = 3.93ms; TI = 900ms; flip angle= 8; 176 slices, FOV= 256mm).

As in a typical Simon task, subjects were presented a box to either the left or right hemifield and were asked to respond to its color. Upon presentation of a red box, subjects were asked to respond with their right index finger, while upon presentation of a green box they were asked to respond with their left index finger (see Figure 1a). Two sessions were conducted per subject, with each session composed of 48 congruent, 48 incongruent trials and 24 null trials of 2.5s. In the null trials, a stimulus was not presented and no response was necessary.

**Figure 1.**
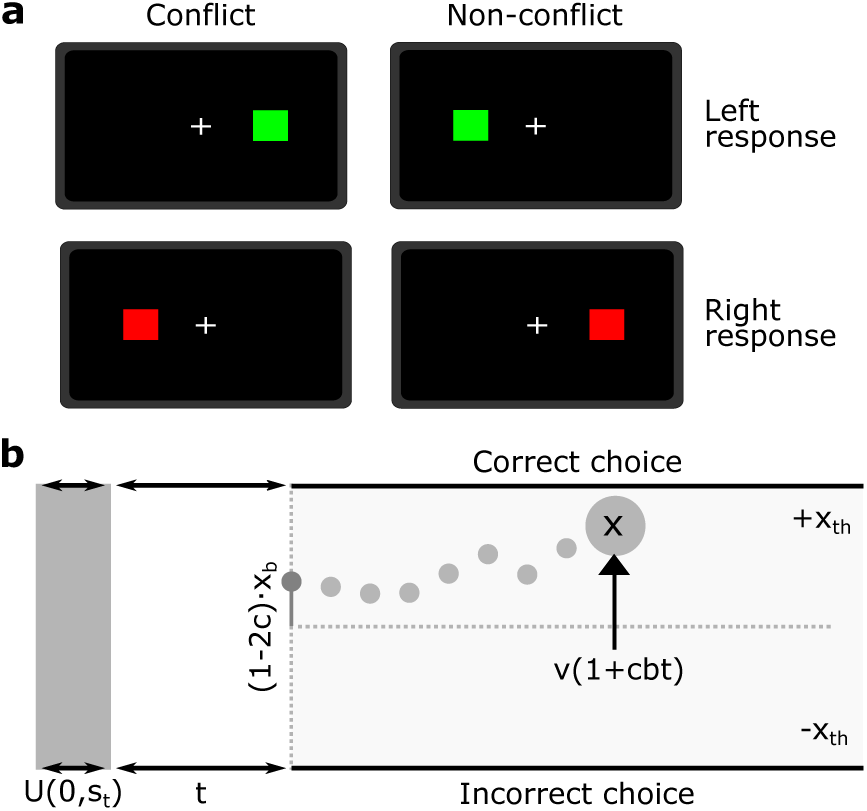
Task structure and model. **(a)** Subjects are presented with a green or red square. A green square indicates that the subject should make a response with their left hand while a red square indicates that the subject should respond with their right hand. Conflict trials are those where the side to which the square is presented to is inconsistent with the required response. **(b)** Decision-variable to bound in the SE-SSM for a non-conflict trial. Parameters are explained in equations 1-3.

### Simon effect model and fitting

#### Model

In previous work (see supplementary material), a SSM was proposed based on the the exact stimulus presentation (composed of visual hemifield, and color) and response (left or right hand). As this information is not present in the available dataset, here we use a mathematically equivalent, accuracy coded form (Figure 1b) where the model is defined in terms of a single stimulus conflict parameter, and consider the response only in terms of whether it was correct or incorrect (eq: 1-3):

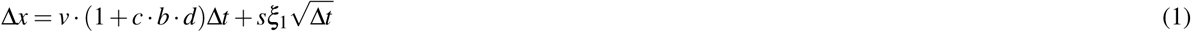

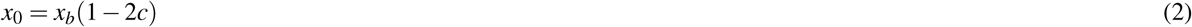

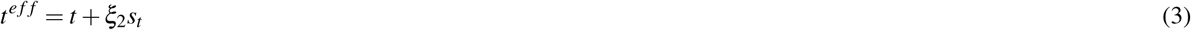

The variable *x* denotes the decision-variable that builds to a threshold *x*_*th*_. The stimulus conflict parameter *c* is set to take a value of 0 when there is no conflict present (i.e. when the stimulus is presented to the visual hemifield matching the required response hand), or 1 otherwise. Parameters *v, s, t* and *x*_0_ correspond to the drift, noise, non-decision time and starting point bias, while *ξ*_1_ represents the Gaussian diffusion process. The effective non-decision time on a given trial *t*^*e f f*^ is defined as the sum of a base non-decision time *t* and a uniform noise process *ξ*_2_ scaled by the noise in the non-decision time term *s*_*t*_. Parameters that are specific to the Simon task are *b*, a supposed conflict counteraction component that enhances the effective drift on conflict trials as its duration *d* (measured in seconds) increases, and *x*_*b*_ a parameter that captures the initial response bias caused by the Simon effect.

#### Basic model fits

Individual subject models were fit by using our SSM fitting framework (see supplementary material) which minimizes the negative log-likelihood by using Matlab’s generalized pattern search algorithm^25^. In order to generate the probability density for a specific set of parameters and conditions we resorted to encoding the probabilistic model decision-variable updates over time into a transition matrix (a similar method was also employed by Brunton et al. 2013^26^). We relied on such a method so that we could make use of likelihoods for model comparison, as well as it being relatively faster to evaluate (and consequently fit) than methods requiring the full simulation of individual decision-variable traces.

The transition matrix is fixed on non-conflict trials, and varies in a time dependent manner in conflict trials because of the *b · d* term. For each probability density evaluation, we discretized an initial decision-variable vector *x* so that it would span 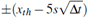, a range large enough to practically avoid being exceeded in a single step of the decision-variable. The *x* entry equivalent to the closest *x*_*b*_ value is set to one. The transition matrix is then iteratively defined for the conflict trials, or initially defined for the non-conflict trials by computing the expected Gaussian distribution for the decision-variable on the following trial for all possible current values of the decision-variable. Entries in the transition matrix outside of the decision thresholds are set to zero, except for where the transition leads to itself (i.e. diagonal entries), in which case they are set to one. The cumulative density function is then calculated by summing the iterated output above and below the model threshold. In order to generate the final density function, the cumulative density is numerically differentiated. We verified that these model fitting procedures worked sufficiently well to recover simulated parameters (see supplementary Figure S3 and S4). Model fits were randomly initialized from the following set of distributions: *v* ∼*G*(3, 1), *x*_*th*_ ∼*G*(0.6, 0.15), *t* ∼*G*(0.3, 0.075), *b* ∼*HN*(2.5), *x*_*b*_ ∼*G*(0.1, 0.06), *s*_*t*_ ∼*U* (0, 0.15). Where, *G*(*x, y*) represents the Gamma distribution parameterized by mean and standard deviation, *HN*(*x*) represents the half Normal distribution parameterized by the standard deviation, and *U* (*a, b*) represents the Uniform distribution parameterized by its two limits. The parameter *s* was fixed to one for all cases.

We initially fitted all our models with the decision-variable resolution set to 0.01, and repeated this five times to confirm that our procedure was consistently discovering global minima. After this procedure, we performed one further pass using the previous best model as a starting point with the decision-variable resolution set to 0.005 for more accurate parameter estimates.

### BOLD processing

#### Pre-processing

Standard processing was implemented in FMRIB’s Software Library (FSL^27^). Separation of non-brain tissue was performed using Brain Extraction Tool (BET^28^) with robust brain center estimation. Functional images were registered linearly to subject’s structural image (BBR algorithm) and non-linearly (FNIRT, 12 DOF, 10mm warp resolution) to a standard Montreal Neurological Institute (MNI) brain template. Functional data was high-pass filtered (0.01 Hz), and spatially smoothed with a 5mm full-width half-maximum Gaussian kernel. The FEAT tool within FSL was used for event-related fMRI analysis. Stimuli were modeled as 100ms long step functions and motion parameters (generated by MCFLIRT^29^) were added to the model as confounds of no interest. FEAT was set to use a double-gamma haemodynamic response function to convolve with our regressors of interest.

#### Model driven across subject BOLD estimates

We hypothesized that if the SE-SSM is describing core components of the Simon effect that we may be able to see correlations between average stimulus activation across subjects and Simon task specific parameters (*b* and *x*_*b*_). To investigate this, our analysis begins by application of a fixed effects general linear model (GLM) with a single set of fixed effect regressors generated by all stimuli for correct responses. This analysis is carried out at first (trial) level and extended to second (session) level, yielding GLM weights, which are then transformed into MNI space. MatlabTFCE^1^ was used to generate p-values corresponding to each voxel after a mask excluding brain-stem and white matter was applied. Specifically for our application, MatlabTFCE calculates the correlation coefficients between GLM weights and parameter values of interest across subjects for each voxel and then applies the Fisher z-transformation. A threshold-free cluster enhancement (TFCE^30^) transformation (default parameters were used: E = 0.5, H = 2) is then applied to these values, and the absolute value is taken (the absolute value is required for the two-tailed test). Generation of voxel *p*-values is then carried out by permutation. The permutation works as follows: at every iteration the z-transformed correlation is calculated from the parameters with respect to shuffled GLM weights, and a maximum z-value across voxels is calculated. For each voxel a count normalized by the total number of permutations performed is incremented every time the maximum z-value exceeded the real z-value, resulting in a maximal statistic permutation family-wise error rate (MSP FWER) corrected two-tailed p-value for every voxel. The p-value map is then split into two dependent on whether the real z-value is positive or negative, in order to generate a positive-tailed p-value map and a negative-tailed p-value map, with the complimentary voxels set to a p-value of 1.

To demonstrate that the model based analysis provides insights not accessible by simply examining the RT, we repeated the same analysis using the average RT across subjects, as well as the average difference in RT between congruent and incongruent trials across subjects.

#### Within trial BOLD estimates

We proceeded to investigate whether patterns of BOLD activation correspond to the development of the expected decision-variable (DV), a method similar in principle to EEG-fMRI analysis used by Philiastides and Sajda^31^. In order to generate the expected decision-variable for each trial, we begin by simulating the fitted SE-SSM. Unlike the methods explained in the model fitting procedure, in this case the decision-variable is explicitly simulated at every time step until a threshold is reached. This process is repeated 750 × 10^3^ times with the full decision-variable trace being saved, together with corresponding RT, and choice (correct or incorrect). For each real trial, we then match all simulated trials with the same choice and RT (up to a resolution of − 25ms), align them to their threshold crossing and average in order to generate the expected decision-variable trace over time. Once the decision-variable trace has been defined for all trials and for all subjects, we extract the average decision-variable in time windows stepping back from the response time. We set the window width to be 50ms, starting centered at 25ms and moved it back in 50ms steps. The expected decision-variable was then extracted from a specific time window, z-scored and fed into a first level GLM which we term ‘DV-GLM’. This GLM additionally takes as its first feature, events corresponding to correct trials, and we carried it through to second level and third level (across subjects) with a mixed effects GLM). Clusters were thresholded at z *>* 2.5 and corrected for multiple comparisons using Gaussian random field theory (GFT) at *p* < 0.05. We repeated this process for each time window, generating an activation map for each.

#### Traditional BOLD analysis

We also conducted a more traditional BOLD analysis where we carry over first and second level GLM analyses to the third level (across subjects) with a mixed effects GLM. Clusters were thresholded at z *>* 2.5 and corrected for multiple comparisons using GFT at *p* < 0.05.

GLMs for this analysis were run with various combinations of variables of interest. A ‘simple GLM’ was initially implemented to test for whether there is a detectable response specific to incongruent stimuli, by using all stimuli as events for a first regressor and only incongruent stimuli as a second regressor. In this analysis, the coefficient matching the second regressor is then indicative of the difference in activation between congruent and incongruent stimuli. In the ‘RT GLM’, we proceeded to investigate whether the stimulus BOLD response is modulated by the RT, by using all stimuli as the first regressor, and the log-transformed and then z-scored RT for the second regressor.

## Results

### Simon effect is present in behavior

The Simon effect is visible in Figure 2a, and we confirmed its presence by performing a Wilcoxon signed-rank test on subject RT averages across conditions (*p* = 0.0006, n = 21, z = 3.42). The average (mean) difference in RT between conflict and non-conflict trials was 25.7ms ±5.7ms (SEM). Standard deviations of the RT were also different between conflict and non-conflict (*p* = 0.0087, n = 21, z = 2.62), a trait common to the Simon effect, and potentially indicative of the involvement of cognitive control. As expected, accuracies in conflict trials were decreased with respect to non-conflict trials although the overall accuracy in both conditions was very high (non-conflict: 97.3% ±0.5% (SEM), conflict: 95.6% ±0.5% (SEM), Wilcoxon signed-rank *p* = 0.0172, n = 21, z = 2.38). The decreased accuracy in conflict trials with respect to non-conflict trials is driven by early responses (Figure 2b-c), an observation which has been incorporated into Simon effect models.

**Figure 2.**
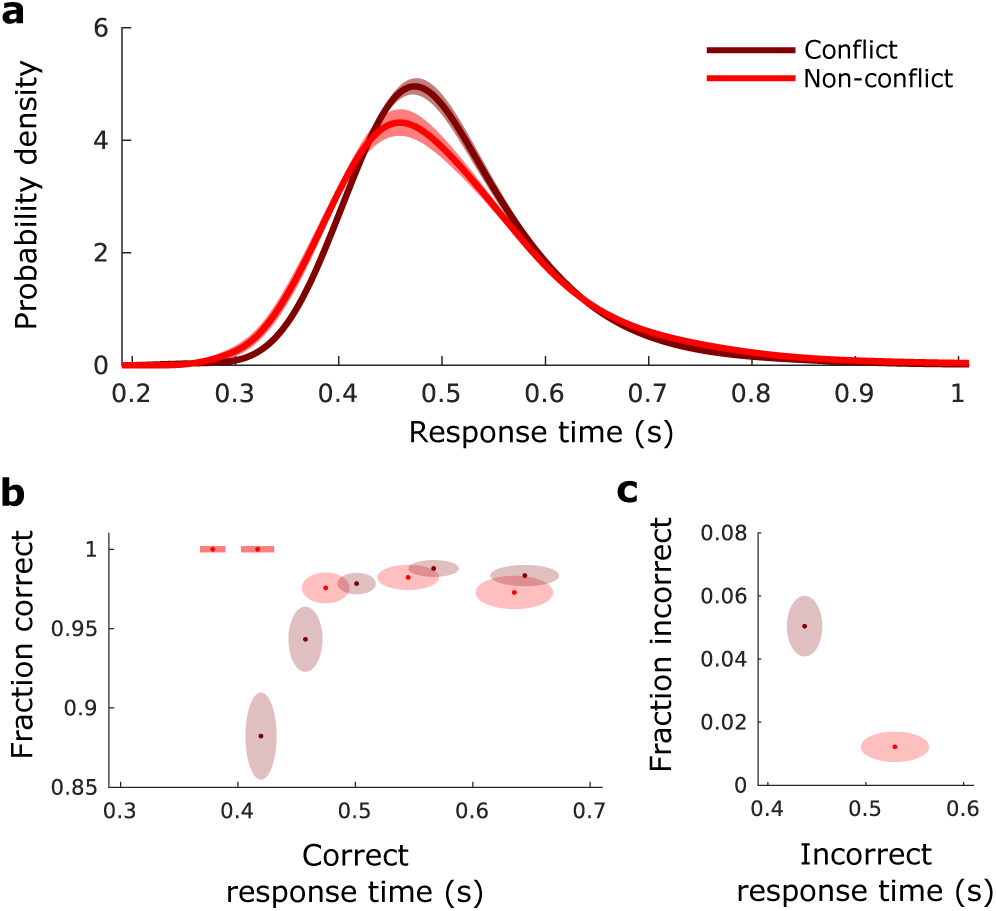
Behavioral data. (**a**) Subject averaged density estimate of correct RT for conflict and non-conflict responses. Simulated data is initially normalized at the individual subject level by dividing by subject standard deviation and then multiplying by grand mean RT, density estimation is performed by convolution with a Gaussian kernel. (**b**) Fraction of correct responses within response time percentile bins, plotted against central percentile RT. Percentiles bin edges were selected to be [0, 20, 40, 60, 80, 100]%. For example, the right most point shows the accuracy for data between the 80th and 100th percentile plotted against the 90th percentile response time. Ellipse width signifies SEM in RT across subjects, while ellipse height signifies standard-error in accuracy across subjects. (**c**) Same as **b** but for incorrect responses, a single percentile of 50% was used due to the low number of incorrect responses.

### Model captures Simon effect

As an informal assessment of the model fit quality for the SE-SSM, we re-sampled from it for each subject, and analyzed the data in the same way that we analyzed the behavioral data (c.f. Figure 2a-c to Figure 3a-c). The model fits demonstrate the Simon effect (Wilcoxon signed-rank test on subject RT averages, *p* = 0.0006, n = 21, z = 3.42). Visual inspection indicates a good match, the main difference being the across-subject variability which has a negligible within-subject component for our model. The average (mean) difference in RT between conflict and non-conflict trials was 29.2ms ± 6.0ms (SEM). Standard deviations of the RT were also different between conflict and non-conflict (Wilcoxon signed-rank, *p* = 0.0046, n = 21, z = 2.83), and accuracies in conflict trials were decreased with respect to non-conflict trials (non-conflict: 98.3% ± 0.4% (SEM), conflict: 96.6% ±0.4% (SEM), Wilcoxon signed-rank *p* = 0.002, n = 21, z = 3.09).

**Figure 3.**
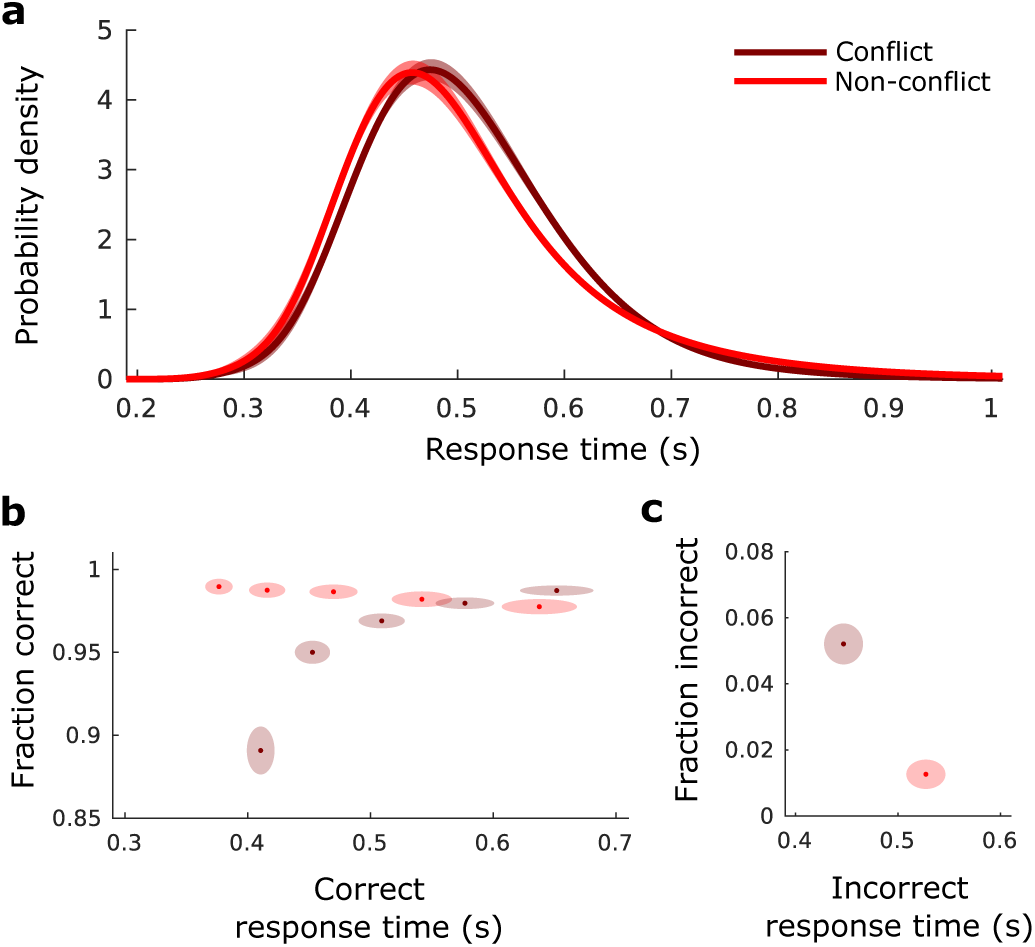
Model fitted simulated data for comparison to Figure 2. Each subject model fit is used to produce 104 responses. (**a**)-(**c**) as in Figure 2.

More formally, we computed Akaike information criterion (AIC) and weighted AIC (w_AIC_^32, 33^) for the SE-SSM (M4), and less complex candidate models. The less complex candidate models considered are: the basic SSM (M1), the basic SSM with the conflict counteraction parameter *b* (M2), the basic SSM with the bias parameter *x*_*b*_ (M3). The w_AIC_ of a given model can be interpreted as the probability of that model being the best model among the candidate set^33^.

We found that candidate models M1 and M2 were not well supported (largest w_AIC_ for 2 subjects each). Models M3 and M4 on the other hand were both well supported, with the largest w_AIC_ coming from M3 for 8 subjects, and M4 for 9 subjects. This data is summarized in Figure 4.

**Figure 4.**
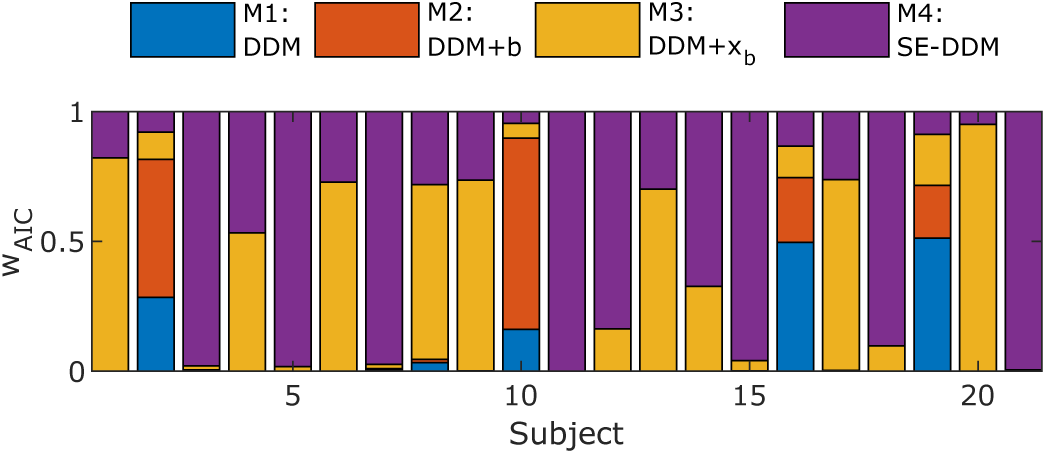
wAIC for the basic SSM (M1), the basic SSM with the conflict counteraction parameter *b* (M2), the basic SSM with the bias parameter *xb* (M3) and the SE-SSM (conflict counteraction parameter *b* and bias parameter *x*_*b*_, M4). M3 and M4 are the best candidate models.

There are several points to note here: unsurprisingly, it seems that the addition of a bias term is essential to capture the Simon effect. On the other hand, the addition of the conflict counteraction term does not always clearly improve the model quality. While the variability between M3 and M4 may be partially due to sampling variability, it seems likely that some subjects simply behave more consistently with M3, while others behave more consistently with M4. Specifically, it should not be a surprise that there is variability in how subjects deploy conflict counteraction during the Simon task, with those that deploy it sparingly being captured best by M3.

Since the SE-SSM (M4) is the best fitting candidate in the largest number of our subjects, and it encompasses all other models (specifically, M3 is a special case of M4 when *b* is close to zero) we proceeded to analyze BOLD activity in terms of this model.

### SE-SSM Across subject analysis

In this analysis we fitted a GLM with a single series of events corresponding to the stimuli. We then computed the correlation between our two task specific SE-SSM parameters (*b, x*_*b*_) and the GLM weight maps across subjects. Activation, as assessed by permutation tests of the TFCE transformed correlations was present in several regions.

We repeated this correlation analysis with average subject RT, as well as the difference in subject RT between congruent and incongruent conditions (ΔRT) and found that no voxels were activated (*p* < 0.05, MSP FWER corrected). This demonstrates a clear benefit of the model based approach: RT and ΔRT are dominated by several sources of variance which are irrelevant to conflict or act to cancel each other out so that direct analysis may be inconclusive, while analysis of the underlying estimates of the contributions shows clear activation.

Correlations with bias *x*_*b*_ across subjects (Figure 5a, orange *p* < 0.05, MSP FWER corrected) were present in multiple regions, most prominently and of particular interest the supplementary motor area (SMA), insular cortex, lingual gyrus, precuneus, posterior and anterior cingulate, the right precentral gyrus as well as the superior frontal and paracingulate gyrus and cerebellum.

**Figure 5.**
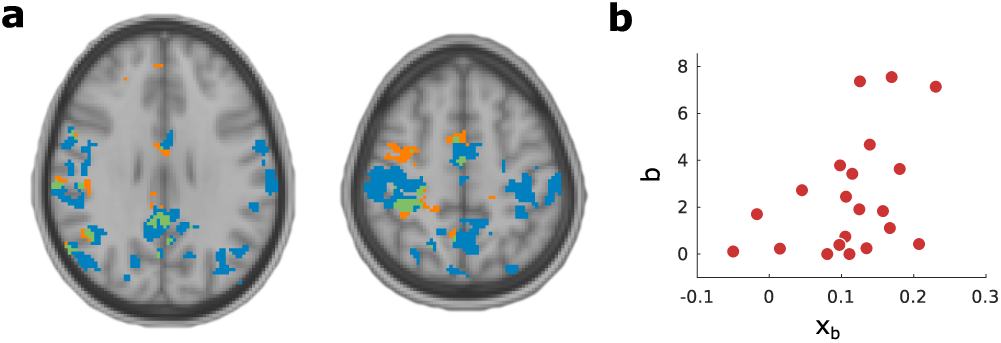
Regions where activity is related to SE-SSM parameters across subjects. (**a**) Regions where activity is related to (*p* < 0.05, MSP FWER corrected) the conflict counteraction parameter *b* are shaded in blue, while regions where activity is related to (*p* < 0.05, MSP FWER corrected) the bias parameter *xb* are shaded in orange. Regions where both parameters highlight significant regions are shaded in green (*p* < 0.05, MSP FWER corrected). (**b**) While regions in **a** are mainly distinct, some overlap may be accounted for by the correlation between the bias and conflict counteraction parameter across subjects (Spearman’s CC = 0.48, *p* = 0.029).

Correlations with conflict counteraction *b* across subjects (Figure 5, blue, *p* < 0.05, MSP FWER corrected) were most prominently present in SMA, insular cortex, lingual gyrus, precuneus, posterior and anterior cingulate, and the left and right precentral gyrus and cerebellum.

The partial overlap in brain activation (Figure 5, green) related to the SE-SSM parameters led us to investigate whether across subjects, the parameters *b* and *x*_*b*_ are correlated. Indeed, as shown in Figure 5b there does appear to be some correlation between the two parameters.

### SE-SSM within trial analysis

Given the RT and whether the choice was correct on a given trial, we estimated the progression of the expected decision-variable (Figure 6). We then used these values averaged over successive 50ms windows using a GLM (see work by Philiastides and Sajda^31^ for a variation of this method applied in EEG-fMRI) in order to extract details regarding the progression of information during a trial. We found clusters of BOLD activity (*p* < 0.05, GFP FWER corrected) with the expected decision-variable extracted from windows centered at −25ms and −175ms. Early activation at −175ms includes the influence of the decision-variable jump, smoothed by the 50ms window, and may represent activation of the automatic pathway^15^ - itself a signature of conflict. It is therefore consistent, that the activated region at this stage is the paracingulate and superior frontal gyrus, neighboring both SMA and ACC. At −25ms activation appears to be dominant in intracalcarine cortex and lingual gyrus.

**Figure 6.**
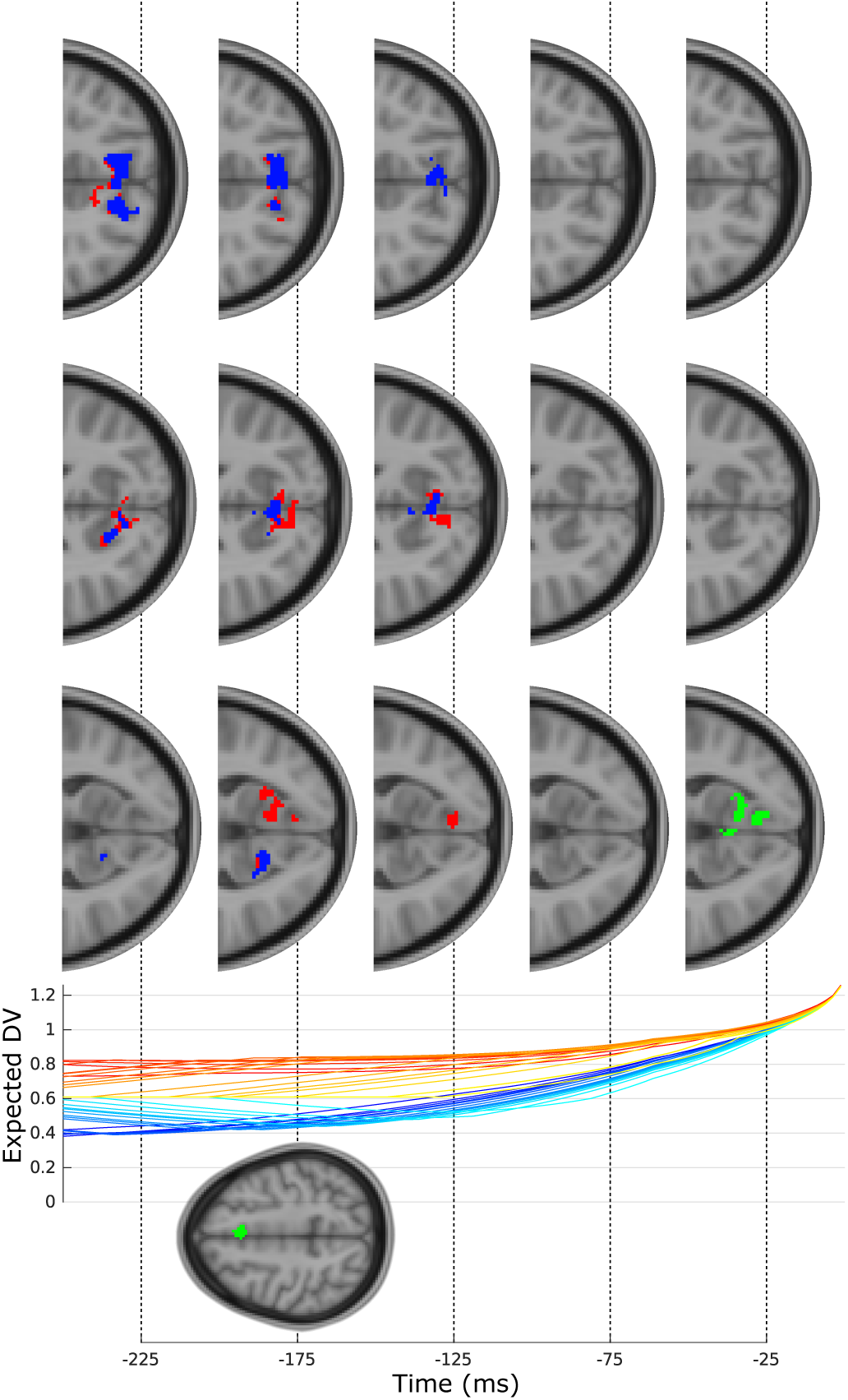
Expected decision-variable BOLD activation relative to response. Top: from left to right, transverse slices showing activation (*p* < 0.05, GFP FWER corrected) from GLM with decision-variable taken from corresponding time. Activation shaded in green stems from GLM using decision-variable pooled from both conditions (we note additional activation at −175ms). Activation shaded in red and blue stem from GLM with the decision-variable split by condition, with red corresponding to a sum contrast, and blue corresponding to conflict only (non-conflict only did not yield any statistically significant regions). Bottom subplot shows an example decision-variable progression for one subject, and an axial slice at −175ms. Orange: non-conflict, blue: conflict, shading dependent on RT rank.

To examine this further, we repeated our DV-GLM but split the expected decision-variable by whether it came from a congruent or incongruent trial. This analysis ensures that only the decision-variable variability within the congruent and, separately within the incongruent conditions can account for BOLD activation, thus removing the direct contribution of conflict. As well as assessing the statistical significance of the two decision-variable regressors, we also examined the contrast of their sum. While the non-conflict decision-variable did not yield any statistically significant BOLD activation, interestingly, both the conflict decision-variable and sum contrast did. The sum contrast activates regions largely overlapping to those activated by the single decision-variable GLM at −25ms, although earlier with respect to the response and extending into cuneal cortex and precuneus.

### Stimulus-response GLM analysis

We also performed a more traditional set of analyses which we termed ‘Simple GLM’ and ‘RT GLM’.

The simple GLM consists of modeling stimulus presentation as a single series of events, with incongruent events as an additional series. The stimulus presentation appears to activate clusters in the visual areas (large sections of occipital and lateral occipital cortex), as well as response related regions such as the SMA, pre- and post central gyrus and the left central opercular cortex. Activation in cerebellum was also present.

Importantly, in contrast, when examining coefficients coding for incongruence relative to stimulus presentation no clusters were present. This suggests (as has been found in previous work^21^) that the activation present due to conflict is subtle and may require more sophisticated analysis as we have presented here.

While incongruent stimuli may not directly lead to clear activation, we considered that RT may in part reflect the process of conflict detection. To test this, we performed a GLM with one event variable related to stimulus as before, and another event variable set to be the z-scored log-RT. The stimulus event clusters for this analysis are broadly similar to those presented for the previous analysis (stimuli and incongruent events, compare Figure 7a to Figure 7b), with the most prominent difference being the lack of a cluster in the left central opercular cortex. The z-scored log-RT event however appears to elicit activity in ACC, as well as bilaterally in insular cortex, and occipital-parietal areas ‘upstream’ to those visible for the stimulus event. These regions include intra/supracalcarine cortex and superior lateral occipital cortex.

**Figure 7.**
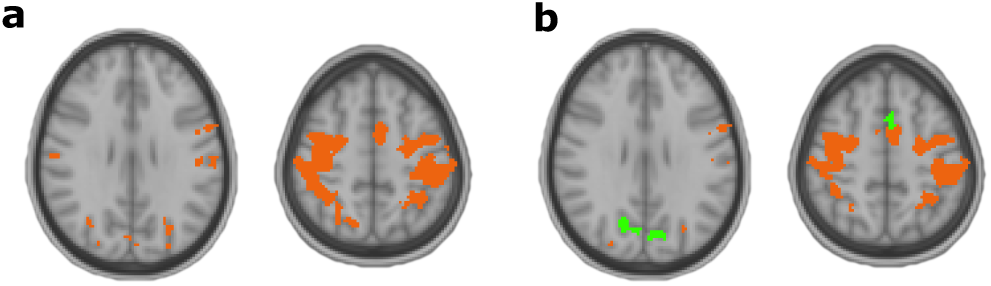
Standard GLM analyses. (**a**) Clusters for simple GLM. Orange clusters represent activation in response to stimulus, and no additional clusters are present in response to an incongruent event. (**b**) Clusters for RT GLM. Orange clusters represent activation in response to stimulus, similar to **a**, while green clusters represent activation dependent on z-scored log-RT events.

## Discussion

In this study, we consider the Simon effect to emerge out of three main components: 1) a representation of the choice relevant stimulus (color), stimulus laterality, and conflict with respect to choice; 2) evidence accumulators for each choice wherein their relative activation is the decision-variable; 3) cognitive control to appropriately distribute the computation across these components. We found that the SE-SSM which incorporates these components yields parameters which correlate with stimulus BOLD activation, while equivalent analysis based on RT does not.

Interpreting the SE-SSM in light of previously developed cognitive models^6, 13, 14^ we suggest that the decision-variable should be considered as equivalent to the relative activation of response accumulators, with the bias term representing the relative strength of the automatic pathway, and the conflict counteraction term linked to attention, monitoring and regulation. Across subject activation may depended on generic traits of the subject and should correspond to global parameter settings that project to regions where an individual trial is being processed, or targets of such regions. We also used the model to analyze the within BOLD correlates of the evolution of the expected decision-variable.

Because the RT distribution of the SE-SSM cannot be written analytically, in previous work we resorted to minimization of the *χ*^2^ statistic^34^ calculated from binned RT data which is generated by direct simulation of the SE-SSM process. Additionally, model comparison was done by examining the *χ*^2^ statistic and application of cross validation, leading to a computationally expensive procedure. In this work, we developed a SSM framework to fit non-standard extensions of the SSM more efficiently. To do this, we encode the transition matrix of the SE-SSM and propagate this forwards in time in order to generate the model likelihoods. Such a method allows for a substantial speed up in model fitting so that we could be confident that we were avoiding local minima by repeating simulations. The method also directly allows us to use existing model comparison techniques without resorting to further simulations. We also note that the model estimation procedure employed, including the calculation of likelihood functions from a non-standard SSM could lend themselves to being incorporated into hierarchical SSM frameworks such as the HDDM^35^.

We found that SE-SSM provided a good explanation for the behavioral data. However, the separation between the SE-SSM and SSM including conflict dependent bias was not as clear as we were expecting. Indeed, when applying the current methods to data from (data not shown^12^), we find that the SE-SSM is strongly preferred over all other candidate models (as was shown via cross-validation). We believe that this inconsistency is likely due to the relative weaker maximum trial time constraints placed on subjects in the current data set, providing less necessity for a strong counter active role of conflict counteraction on bias.

Using a GLM with all events, and additionally incongruent events as regressors, only yielded a statistically significant pattern of activation in response to all events. Performing a similar GLM with RT as opposed to specifically modeling incongruent stimuli yielded a statistically significant ACC cluster. However, it is not clear whether this should be directly related to conflict as the RT ultimately is dependent on a variety of factors. This highlights a potential advantage of the model based approach: decomposition of accuracy and RT into model parameters acts to isolate concepts of conflict counteraction and bias which can then be used to interpret the data.

The ACC which we find to be clearly activated in our across subject analysis has been implicated in conflict monitoring^36^ and resultant specification of a control signal^13^, which we hypothsesise should be related to conflict counteraction. It is also generally found to be activated in several previous Simon effect fMRI studies^18–20, 37^, and is known to be activated with insular cortex^38^. Aside from sensorimotor areas, other clear regions of activation are precuneus and the posterior cingulate. In previous work, we ascribed the concept of attention to our conflict counteraction model parameter *b*, and so it is interesting to note that precuneus has also been found to be active in previous Simon effect studies, and has been implicated in shifting attention to different spatial locations^39^. However, we currently make no strong claims regarding specific theories of attention^40^.

While activation related to the bias term was present in frontal and paracingulate giri, we found that its activation largely overlapped with the activation of the conflict counteraction parameter. This would be partially predicted by the correlation between conflict counteraction and bias which is in itself unsurprising: subjects with a strong bias, may compensate by enhancing their conflict counteraction. This study, and previous work suggests that two task specific parameters are required to fully capture behavior in the Simon task, so we hypothesize that the overlap of the brain activity is due to the interwoven and distributed nature of the decision making process. We cannot however rule out that there is a more parsimonious model structure that would allow the Simon effect to be captured with a single variable which itself optimally reflects brain activity. This question will need to be resolved in future studies where the conflict counteraction parameter is modulated experimentally.

Within a trial, the consistent activation of a single region shows a location where a decision-variable could be directly represented, or where components required for its calculation are represented. For example, regions activating during the early representation of the decision-variable progression could be related to automatic activation, while later activation could be related to evidence integration and conflict counteraction. An example of our interpretation for the specific process that is occurring during a Simon effect trial is as follows: a left indicating stimulus is presented to the right hemifield. An automatic pathway is activated, and feeds into a response accumulator. This pathway is quickly inhibited indirectly, or its activity is inherently transient so that while an impact has been made on the current state of the accumulators, further impact is reduced. We model this as a delta function at a moment corresponding to the non-decision time - although a more natural process starting somewhat earlier is likely to occur in reality. The non-automatic, task specific process is activated less quickly - and begins to feed into the pre-motor accumulators once the automatic pathway is no longer having a novel impact. On some occasions the inhibition is not fully deployed, or noise in the decision process acts to initiate an action at this early stage. Usually however, this is not the case and evidence from the non-automatic pathway gradually pushes the correct response accumulator towards its threshold. Due to time constraints of the task, or the inherent cost of time however, a compensation mechanism takes hold to reduce the influence of the initial bias. Such a mechanism should make use of the representation of the decision-variable in terms of the initial stimulus laterality: a measure of conflict which changes as the trial progresses. Monitoring of this conflict could in turn be used to accelerate the decision-variable towards a bound.

In our within trial analysis, we attempt to examine the progression of the decision with high temporal resolution despite fMRI’s low temporal resolution using a method inspired by EEG-fMRI analysis^31^ and the concept of examining the expected decision-variable^41^. We found clear representation of the decision-variable early and late relative to the response. Early activation (175ms prior to response) appears to be in paracingulate and superior frontal gyri which neighbors SMA and ACC. It seems possible that this corresponds to early detection of conflict, or deployment of conflict counteraction in response to conflict, which would be represented in the decision-variable at an early stage in a graded manner due to the smoothing effect of our temporal windows. We also detect late activation relative to response time (−25ms) of intracalcarine cortex. This type of activation would be consistent with full representation of a decision-variable like feature prior to response, although its presence in early visual areas late in the decision is surprising. We were curious as to whether we might find a separate representation of the conflict specific decision-variable, and found that unlike the decision-variable taken from all trials, activation is more strongly represented in precuneus. Consistent with previous studies, we hypothesize that the posterior medial frontal cortex coordinates with precuneus to monitor the level of conflict, and deploy conflict counteraction for the purposes of modulating the automatic pathway.

A weakness of the current study, particularly relevant for the time resolved analysis stems from the data being cast in terms of conflict and accuracy, rather than stimulus and response: stimulus and response laterality influence the decision-variable in different ways than conflict and accuracy producing potentially different patterns of activation. For example, we would expect the early stimulus-response decision-variable to be lateralized in the occipital region, while the late decision-variable may be lateralized in the motor or pre-motor region. In this study, we detect early ACC activation in our within-trial analysis, potentially corresponding to conflict detection. While it may be the case that conflict detection is occurring early in the trial, an alternative explanation is that early expected decision-variable is just an indirect signature of conflict (much like RT is), which is itself only represented later in a trial. This possibility can be analyzed more carefully with exact stimulus and choice information, where conflict is not directly confounded with the early decision-variable. In future work, we intend to address these points, and note that improved experimental conditions may also allow us to analyze the across trial intermediate temporal domain and influence of conflict history in the context of the proposed model (see supplementary material Figure S1 and S2).

## Acknowledgements

We thank Anna Jasper and Tao Tu for comments. This work was partially funded by the Army Research Laboratory (ARL) under cooperative agreement number W911NF-10-2-0022, we acknowledge computing resources from Columbia University’s Shared Research Computing Facility project, which is supported by NIH Research Facility Improvement Grant 1G20RR030893-01, and associated funds from the New York State Empire State Development, Division of Science Technology and Innovation (NYSTAR) Contract C090171, both awarded April 15, 2010. We acknowledge the OpenfMRI project and NSF Grant OCI-1131441 (R. Poldrack, PI) for making this data accessible. Declarations of interest: none.

## Data availability

The datasets analyzed during the current study are available from the OpenfMRI database under an ODC Public Domain Dedication and Licence (PDDL). Its accession number is ds000101^24^.

## Author contributions statement

J.R.M. conceived and performed analysis. J.R.M and P.S. wrote manuscript. All authors reviewed the manuscript.

## Supplementary Material

### Full model

In previous work we defined a model to capture the Simon task which we reproduce here with parameters consistent with the present manuscript:

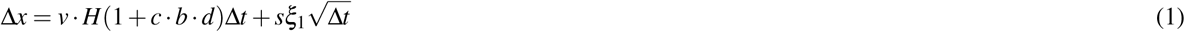

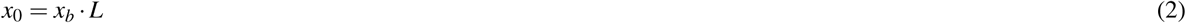

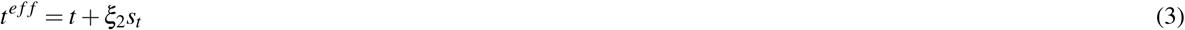

Where the conflict 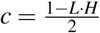, and *H* and *L* are trial dependent variables with values of 1 or +1 corresponding to the required handedness of the response (color), and visual hemifield of the stimulus respectively. All other parameters are defined as presented in the main text, although we would also like to note that we refer to the parameter *b* as conflict counteraction as opposed to attention.

The model used in this manuscript removes the explicit dependence on *L* and *H*, and is instead written in terms of the presence of conflict.

### Additional analyses

We pursued two additional analyses which we include here for completeness.

#### Across trial BOLD estimates

We were interested in estimating model parameter fluctuations across trials, which we consider to be likely related to learning processes, stimulus strength, as well as random fluctuations in performance. Regions involved in adaptation across trials (modeled as changes in SE-SSM) should either directly correspond to regions that are activated within trial (for example, they may be changes in recurrent excitation in decision circuits) or project to these regions. Examples of such regions are SMA and lPFC, which may also be involved in conflict counteraction and the specification of control signals in response to conflict. While stimulus strength is fixed throughout the experiment, noise in its cortical representation may be present, and this would conceivably manifest in visual areas.

The methods we have introduced thus far enable us to look at brain regions that appear related to our parameters across subjects, and to search for activity correlated with the expected decision-variable at different times within a trial. In order to examine the temporal domain between these, we investigated whether average deconvolved activity in individual voxels could be used to augment the SE-SSM on a trial by trial basis. Deconvolution was carried out using the ‘least squares - separate’ (LS-S) method^1^. For our purposes, this involves repeating GLM fits for each trial, where for each model fit we assign one regressor to a single event matching the stimulus presentation on the trial of interest, and another regressor to absorb all other stimulus events. The regression parameter corresponding to our trial of interest is then treated as the deconvolved activation for this specific trial.

For this analysis, we considered several extensions to the base SE-SSM, for example the augmentation of the drift parameter in a trial dependent manner:

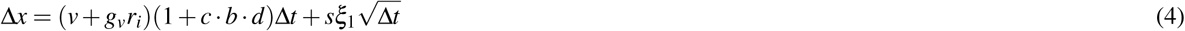

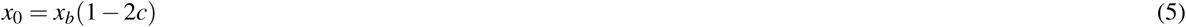

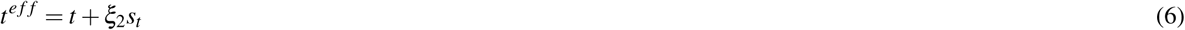

Where in equations 4-6, *r*_*i*_ is a value extracted from trial *i* of an fMRI derived measure. For example, *r*_*i*_ may be the average activation of deconvolved BOLD activation in a specific voxel for trial *i*. The parameter *g*_*v*_ acts to scale the individual trial contributions *r*_*i*_ of these fMRI derived measures.

Two computational difficulties arise when attempting to estimate individual trial parameters based on individual trial voxel activation. Firstly, due to the individual trial modulations, the model likelihood is now slightly modified on each trial, and consequently has to be computed for each trial as opposed to each condition. This is problematic as the time to calculate the likelihood for a set of parameters then scales with the number of trials. Secondly, if a whole brain analysis is required, then this model fit has to be performed at each voxel, which is also impractical. To address these two problems, we make the following compromise: we fix the model parameters at their final estimate (as described in the main text), and then introduce the scaling parameter for *r*_*i*_ (i.e. *g*_*v*_ in the example above). We then estimate the probability density function (dependent on RT, conflict, correctness) for different linearly scaled, zero centered propositions of the impact of the scaling parameter (i.e. *g*_*v*_*r*_*i*_ propositions in the example above). The densities are then made independent of RT, conflict and response by conditioning on the subject response data. Finally, for each voxel we attempt to estimate the parameter *g*_*v*_ by indexing into our candidate density functions with *g*_*v*_*r*_*i*_ and maximizing the sum of the log-likelihood over trials.

After conducting this analysis for each subject, we can then perform a TFCE permutation analysis, although rather than calculating z-transformed correlations (as in the main text) we calculate the z-statistics directly, and compare to permutations generated by randomizing the sign of the parameter across subjects.

We repeated this analysis for individual trial modulation of the bias parameter *x*_*b*_, the conflict counteraction parameter *b*, the model threshold *x*_*th*_, and the drift parameter *v*.

While we found no clear coupling for bias and threshold, both conflict counteraction and drift result in substantial activation across cortical regions.

Small changes in drift and conflict counteraction both lead to similar changes in RT, and it is therefore not possible to say whether one parameter is playing a role while the other is not. Additionally, conflict counteraction is associated with less data (because its effect is zero on non-conflict trials) so a direct comparison is difficult to interpret. Having said this, we repeated this analysis while optimizing drift and conflict counteraction simultaneously and found that no conflict counteraction clusters remained significant. We take this as an indication that the trial-to-trial fluctuations are dominated by changes in drift. Particular regions of interest that are activated in this analysis are cingulate, cuneal, precuneal and posterior cingulate, as well as Superior temporal gyrus (STG), Inferior frontal gyrus (IFG), Medial frontal gyrus (MFG) and the inferior parietal lobule.

In this analysis, regions involved in adaptation across trials modeled as changes in SE-SSM parameters should either directly correspond to regions that are activated within trial (e.g. they are changes in recurrent excitation in decision circuits) or project to these regions. We were expecting pre-SMA activation as well as DLPFC activation, and potentially dACC. We found that these were indeed activated, although other activation was also present. Other activated regions may simply represent trial-to-trial differences in stimulus processing which are irrelevant to the Simon task. It is also worth noting that on an individual trial, changes in drift and changes in conflict counteraction lead to similar changes in RT, making it hard to attribute specific regions to particular processes.

**Figure S 1.**
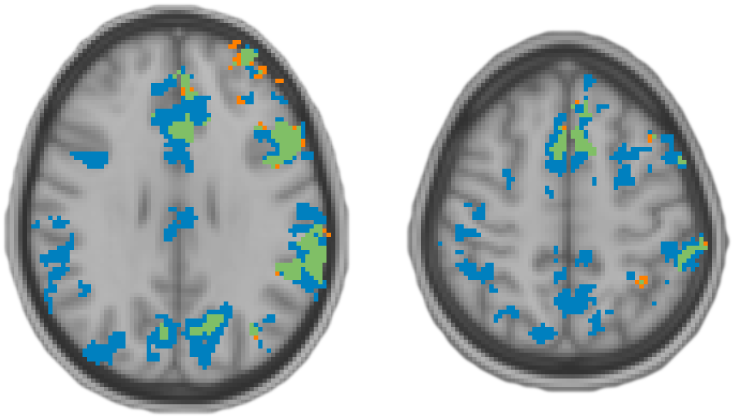
Across trial parameter fluctuation analysis. Blue: activation (*p* < 0.05, GFP FWER corrected) related to the drift term, orange: related to the conflict counteraction parameter, green: related to both conflict counteraction and drift parameters.

#### History model

It is known that presence of conflict influences RT in subsequent trials^2, 3^. We therefore extended the SE-SSM with history terms:

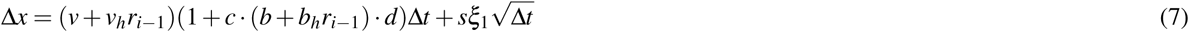

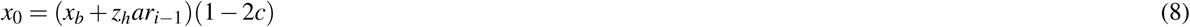

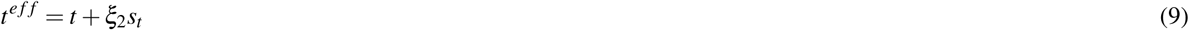

Where *r*_*i*_ = *c*_*i*_ − 0.5 represents the presence of conflict on the previous trial, and *v*_*h*_, *b*_*h*_ and *z*_*h*_ represent parameters to be fitted. We initially fitted each candidate history parameter independent of the other, however to exclude the possibility that both *z*_*h*_ and *b*_*h*_ are needed for the fit, we additionally ran this combination together.

We found that including history into our model, indeed improved the overall model fit quality as shown in 2a. In particular, the model including *z*_*h*_ was favored over including *b*_*h*_ and *v*_*h*_ as well as over the SE-SSM. We wondered whether *z*_*h*_ and *b*_*h*_ together in the same model might improve the fit quality further, but found this not to be the case.

Given that the fitted *z*_*h*_ are on average negative for our subjects, the interpretation is that when a previous trial was a conflict trial, a mechanism to reduce the bias induced by conflict takes hold for the following trial. This is in contrast to recruitment of conflict counteraction which occurs within the same trial.

In order to look for neural correlates of history, we repeated our SE-SSM across subject TFCE based analysis with the parameter *z*_*h*_. However, we found no statistically significant (*p* < 0.05, MST FWER corrected) activation. As an exploratory step, we also investigated whether the *b*_*h*_ parameter was correlated, and indeed found some regions of activation although not in ACC or DLPFC where they might have been expected^3^. We believe that the activation found in this analysis however, is likely due to the general correlation between *b*_*h*_ and *b* itself (Figure 2c) so we do not attempt to interpret this further.

**Figure S 2.**
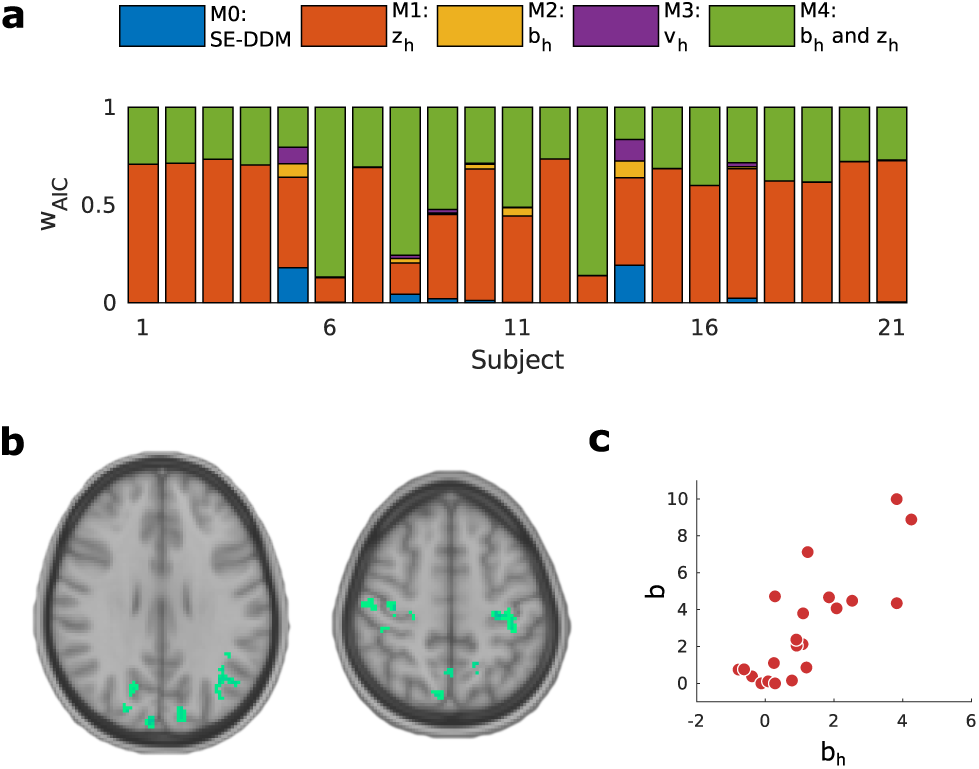
History model, evaluation and results. (**a**) Model incorporating conflict history in the bias term has the highest w_AIC_, improving over the SE-SSM, and being preferable to alternative history configurations. (**b**) While the *z*_*h*_ parameter does not seem to be related to brain activity when investigated with our across subject analysis, the parameter which is shown here in green *b*_*h*_ does. (**c**) However this is likely due to the strong correlation between *b*_*h*_ and *b* (Spearman’s CC = 0.76, *p* = 1 × 10^−4^).

### Model fitting toolbox

In order to fit the SE-SSM, as well as the various nested models and extensions provided, we developed an Matlab toolbox^1^. For parameter initialization, this toolbox incorporates some core functions from the HDDM toolbox^4^ translated from Python to Matlab, as the speed of these methods^5^ is unmatched for the SSM. However, in order to generate density estimates with which to calculate likelihoods for our non-standard SSM candidates, we resorted to encoding the probability distributions of the decision-variable given the value into a transition matrix. The toolbox allows for rapid modification and testing of new models by inheriting from the base SSM class, and then redefining the density estimation.

### Model fitting evaluation

In the ideal case where the underlying behavioral data really comes from a process which is captured by our model, we sought to confirm that our parameters could be recovered with sufficient confidence.

In order to do this, we took our previously fitted parameter values and sampled from each subject the number of trials that they performed. To replicate the procedure described in the manuscript, we repeated this fit with random starts five times. We also repeated this procedure three times to gauge the variability induced by the limited number of samples available for each subject.

We found that as shown in Figure 3, the parameters are well recovered, with correlations for the parameters between true and recovered model being generally greater than 0.8. The parameter that is recovered least well is the conflict counteraction parameter *b*, as expected since this is effectively operating with half the trials since it is only relevant in the presence of conflict by our model definition.

**Figure S 3.**
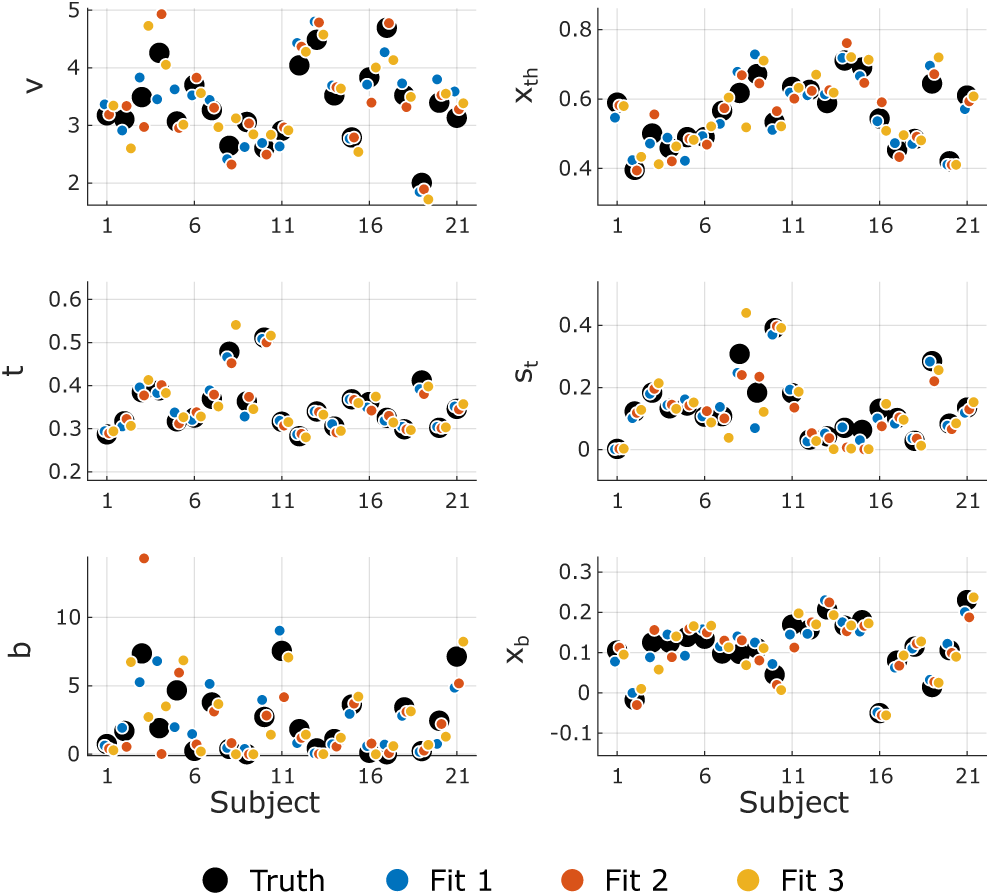
Model fitting parameter recovery. Black circles mark the original underlying parameter fits. Yellow, orange and blue circles mark recovered parameters, with the colors indicating the different samples. Multiple presentations of identically colored circles indicate repeated fit attempts with different random starts.

**Figure S 4.**
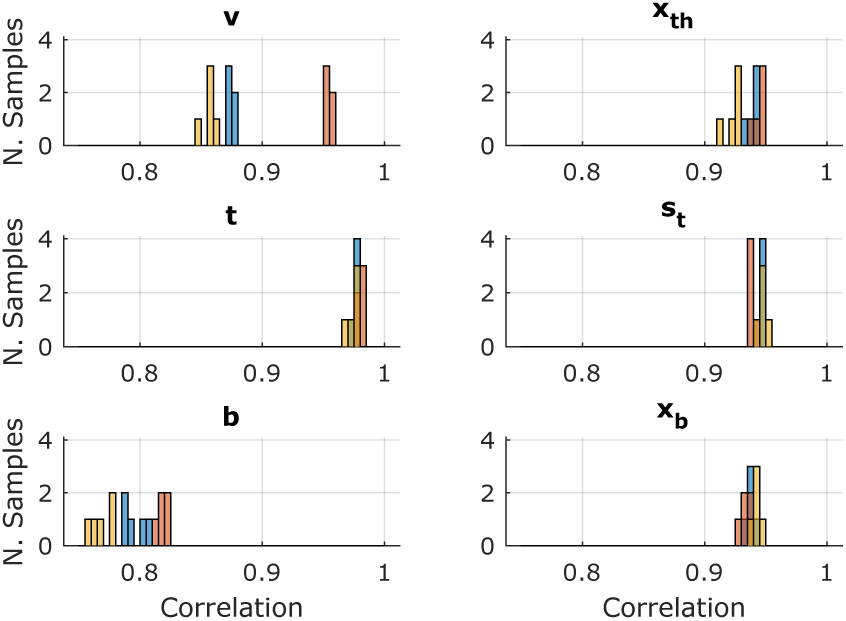
Model fitting recovery correlation of original parameters with fitted parameters. Each color (see Figure 3) represents a different sample from the original model, with each correlation value stemming from a comparison between the original model parameter and the recovered parameters initiated with a different random start.

## Footnotes

1 https://github.com/jrmxn/malleable-ssm

1 https://github.com/markallenthornton/MatlabTFCE

